# Model-Informed Unsupervised Deep Learning Approaches to Frequency and Phase Correction of MRS Signals

**DOI:** 10.1101/2022.06.28.497332

**Authors:** Amir M Shamaei, Jana Starcukova, Iveta Pavlova, Zenon Starcuk

**Author notes:** **Corresponding Author:** Amir M Shamaei, Magnetic Resonance Group, Institute of Scientific Instruments of the Czech Academy of Sciences, Kralovopolska 147, CZ 61264, Brno, Czech Republic. **Code:** https://github.com/amirshamaei/DeepFPC.

## Abstract

**Purpose:** A supervised deep learning (DL) approach for frequency-and-phase Correction (FPC) of MR spectroscopy (MRS) data recently showed encouraging results, but obtaining transients with labels for supervised learning is challenging. This work investigates the feasibility and efficiency of unsupervised DL-based FPC.

**Method:** Two novel DL-based FPC methods (deep learning-based Cr referencing [dCrR] and deep learning-based spectral registration [dSR]) which use a priori physics domain knowledge are presented. The proposed networks were trained, validated, and evaluated using simulated, phantom, and publicly accessible in-vivo MEGA-edited MRS data. The performance of our proposed FPC methods was compared to other generally used FPC methods, in terms of precision and time efficiency. A new measure was proposed in this study to evaluate the FPC method performance. The ability of each of our methods to carry out FPC at varying SNR levels was evaluated. A Monte Carlo (MC) study was carried out to investigate the performance of our proposed methods.

**Result:** The validation using low-SNR manipulated simulated data demonstrated that the proposed methods could perform FPC comparably to other methods. The evaluation showed that the dCrR method achieved the highest performance in phantom data. The applicability of the proposed method for FPC of GABA-edited in-vivo MRS data was demonstrated. Our proposed networks have the potential to reduce computation time significantly.

**Conclusion:** The proposed physics-informed deep neural networks trained in an unsupervised manner with complex data can offer efficient FPC of MRS data in a shorter time.

## Introduction

In Magnetic Resonance Spectroscopy (MRS), typically, more transients are acquired and averaged to increase the low signal-to-noise-ratio (SNR) ^1^. However, individual transients might have different frequency and phase shifts because of hardware imperfections, physiologic processes, or other instabilities ^2,3^. Averaging transients without Frequency and Phase Correction (FPC) would result in line-broadening and lineshape imperfection of the combined MRS signal. Thus FPC should be performed for each transient before averaging. It is even more critical to use accurate FPC while utilizing spectral-edited MRS ^1^ methods to prevent artifacts caused by subtraction. Thus, FPC is a consensus-recommended and effective step ^4^ in MRS signal processing.

Several FPC approaches have been developed ^3,5-10^. FPC methods can be classified into absolute and relative methods ^7^. Absolute approaches correct each individual transient absolutely, whereas relative methods align the transients to a reference signal. A commonly used approach for FPC is to use the water peak and read the phase and frequency from it ^6,8,11^. Another approach is to fit a certain metabolite peak to a model ^12^ and then estimate the frequency and phase shifts from the model. One approach that has been proposed and evolved recently is spectral registration (SR) ^3,5,10,13^. SR fits each signal to a reference signal in the time domain through the adjustment of frequency and phase terms. Even though SR works very well for small shifts, it struggles with larger shifts and signals with low SNR. Modified versions of SR successfully addressed some of the mentioned problems ^5,10,13^. Most of these approaches are time-consuming for large datasets, such as high-resolution MRSI datasets, which may have thousands of spectra.

The recent success of Deep Learning (DL), one of the latest machine learning (ML) approaches, in a wide range of tasks, including the MR field ^14,15^, suggests that it could also handle FPC. Recently, DL-based solutions have been proposed for metabolite quantification in the frequency domain ^16,17^, detecting and removing ghosting artifacts ^18^, FID reconstruction ^19^, automatic peak picking ^20^, enhancement of MRSI spatial resolution ^21^, and identifying and filtering out poor quality spectra ^22^. It has been shown that DL can also be employed for FPC ^7,23^ and could speed up FPC once it has been successfully trained. This approach, using two separate networks in sequence to estimate frequency and phase, showed encouraging results. The first network was trained for frequency shift estimation using the magnitude of frequency- and phase-shifted spectrum as the input and the known frequency shift as the output. Subsequently, the second network was trained for phase shift estimation using real parts of the frequency-corrected spectrum as the input and phase shift as the output. In this approach, any error in the first step (frequency correction) may bias the phase shift estimation. Training two networks is a computationally expensive task. Moreover, the networks were trained in a supervised manner using simulated data. Any discrepancy between the in-vivo and the simulated spectra may result in errors in frequency and phase shift estimation. The true output values are unknown in MRS data, and obtaining hundreds of spectra with labeled frequency and phase shifts is almost infeasible. Unsupervised learning may eliminate the drawbacks of supervised learning.

FPC is traditionally described by adjusting two parameters (frequency and phase). Therefore, it is natural to expect that the variability of all the signals in the set acquired for SNR improvement should have a very low-dimensional representation. One of the methods for non-linear dimensionality reduction is manifold learning, which assumes that the available high-dimensional data vectors are embedded in low-dimensional manifolds ^24,25^. These low-dimensional manifolds can be learned by deep autoencoders (DAEs)^26^, which automates feature extraction by merging all relevant into a cohesive framework. A DAE with a common architecture ^14,26^ is not able to learn to estimate the frequency and the phase shift of a transient since the features in a low-dimensional space might not be readily interpretable. Therefore, a DAE can be redesigned to have two functions: a function for non-linear mapping between the input and certain features (frequency and phase shifts) in a two-dimensional space, and another function, for reconstructing the input from those features. Accordingly, we designed a DAE network that can learn in an unsupervised manner to estimate the frequency and phase shifts of MRS data. The proposed method takes advantage of the parametric analytical approach and embeds it into the DAE to estimate the frequency and phase shifts of a transient.

The proposed network was trained and validated using a simulated dataset in which ground-truth knowledge was available and evaluated using phantom and in-vivo MEGA-edited MRS data obtained from the publicly accessible Big GABA repository ^27,28^. The FPC performance of our proposed network was compared with the commonly used FPC methods; namely, SR ^3^, spectral registration over a limited frequency range (SRF) ^3^, frequency domain correlation (Corr) ^5^, frequency domain correlation over a limited frequency range (CorrF) ^5^, and Creatine referencing (CrR) ^12,29^, in terms of precision and time efficiency.

## Methods

### 1.1 Data Normalization

In contrast to the application of DL in machine vision or speech recognition, where the input data can be normalized by a non-linear transform, MRS signals must be normalized by a linear transform. In this study, each complex signal *S*(*t*) was rescaled as

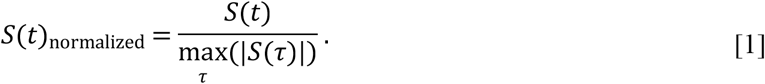

A similar approach could be dividing the signal by the absolute value of the first point of the signal, but it would be less generally applicable because the initial point can be influenced by the filtering processes in the receive chain ^30^, or the maximum may occur later in an echo or with coupled resonances.

### 1.2 Data Augmentation

It is known that the sufficient size and diversity of data are important factors for the effectiveness of most DL models ^14^. However, having rich and sufficient datasets is rare ^31^ in the field of MRS and MRSI. Data augmentation is a viable option, which simulates credible data by minor alterations to data in a small existing dataset. Data augmentation used in computer vision applications to reduce the generalization error of models ^14^ applies flips, translations, and rotations ^14^, which would be meaningless in spectroscopy. To generate credibly varied FID signals, we chose a set of physics-informed alterations, simulating the practical data variability:

1. Frequency shift
2. Phase shift
3. Apodization (line broadening)
4. Amplitude change
5. Adding noise
6. Adding a nuisance peak (residual water and lipids)

### 1.3 Data Sets

#### 1.3.1 Simulated Dataset

A simulated dataset was used to employ in-silico ground truth information for evaluating the performance of our proposed networks and for comparison with the commonly used FPC methods. The simulated dataset was obtained by alternating a single MR signal acquired from a rat brain as described below.

A single-voxel spectroscopy (SVS) MR in vivo signal was acquired from a rat s right hippocampus (256 transients, voxel size = 1.5 × 1.5 × 4 mm^3^) in a 9.4 T small animal MR system (Bruker BioSpin MRI, Ettlingen, Germany) using a PRESS sequence (spectral width = 4400 Hz, 2048 points, TE = 16.5 ms, TR = 2500 ms) with water and outer-volume suppression by VAPOR ^32^. The signal (further referred to as the basis signal *S*_basis_(*t*)) was created after transients were corrected for *B*_0_ instability due to eddy currents as well as *B*_0_ drift, and averaged using Bruker proprietary software, Paravision.

The simulated dataset, containing 24000 artificial signals, was generated from *S*_basis_(*t*) by an augmentation procedure. The basis signal was multiplied by factors drawn from a normal distribution with a mean of 1 and a standard deviation of 0.1. Then a set of lipid and a set of unstable residual water nuisance peaks, generated using Eq. 2 (parameters are listed in Table 1), were added to signals, randomly and independently.

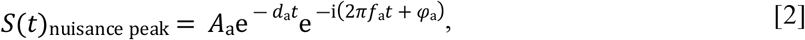

where 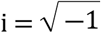, *t* is a vector of time points, and *A*_a_, *d*_a_, *f*_a_ and *φ*_a_ are the amplitude, the damping factor, the precession frequency, and the phases of the nuisance peak, respectively. Signals containing the lipid peak, the unstable residual water peak, and both peaks were labeled as LC, UW, and UW & LC, respectively.

**Table 1.** Parameters for generating the simulated nuisance peak. A_a_, *d*_a_, *f*_a_ and *φ*_a_ are the amplitude, the damping factor, the precession frequency, and the phases of the nuisance peaks, respectively. The spectrometer frequency *f*_0_ = 400 MHz.

| | $A_a$ | $d_a \text{ (s}^{-1}\text{)}$ | $f_a/f_0 \text{ (ppm)}$ | $\varphi_a \text{ (rad)}$ | Number of signals |
| --- | --- | --- | --- | --- | --- |
| **Unstable residual water (UW)** | $\mathcal{N}(1,0.1)$ | $\mathcal{N}(100,10)$ | $\mathcal{N}(0,0.02)$ | 0 | 12000 |
| **Lipid (LC)** | $\mathcal{N}(20,0.1)$ | $\mathcal{N}(300,30)$ | $\mathcal{N}(1.2,0.07)$ | $\pi/2$ | 12000 |

All artificial signals were further apodized by normally distributed random dampings corresponding to Lorentzian linewidths with a mean of 2 Hz and a standard deviation of 0.2 Hz. Then, uniformly distributed artificial frequency and phase offsets in the range of −20 to 20 Hz and −90° to 90°, respectively, were applied to signals. Before the normalizing step, the SNR of signals (time origin magnitude to noise standard deviation) was set in the range of ∼9 to ∼27 by introducing random complex Gaussian white noise. Signals in the simulated dataset were shuffled randomly, and 90% of the dataset was allocated to the training subset, 9% for the validation subset, and the rest 1% for the test subset.

#### 1.3.2 Phantom Dataset

Phantom data was used to assess the performance of our methods in the absence of a large number of training data measured during temperature-dependent changes in the *B*_0_ field.

A SVS MR signal was acquired in a phantom of known metabolite concentrations (NAA 17.5 mmol/L, Glu 26.2 mmol/L, mIns 17.5 mmol/L, Cr 12.7 mmol/L, Tau 10.1 mmol/L; 2048 transients, voxel size = 3 × 3 × 3 mm^3^) in a 9.4 T small animal MR system (Bruker BioSpin MRI, Ettlingen, Germany), while the temperature of the phantom was altered between 35 °C to 40 °C and the frequency adjustment of the scanner was switched off, using PRESS sequence (spectral width = 4400 Hz, 2048 points, TE = 16.5 ms, TR = 2500 ms) with water and outer-volume suppression by VAPOR ^32^.

The phantom dataset, containing 24000 artificial signals, was generated as follows: We selected 400 (out of 2048) of the acquired transients as basis signals, randomly; then, a subset of 60 signals was generated from each basis signal by the same augmentation procedure as used for the simulated dataset except that the basis signal was not multiplied by factors, the nuisance peaks were not added, and the SNR of signals was set in the range of ∼7 to ∼70. Finally, all subsets were stacked together to create the final training dataset. The rest of the 1648 transients were used as an unseen test subset.

#### 1.3.3 Big GABA Dataset

Data from the public repository Big GABA ^27,28^ were used to demonstrate the applicability of the proposed method on FPC of edited in-vivo signals. We selected 48 GABA-edited MEGA-PRESS subsets (subjects) acquired on Siemens scanners from 4 different sites (S1, S5, S6, and S8, 3 Tesla field strength, spectral width = 4000 Hz, 4096 points, TE = 68 ms; ON/OFF editing pulses = 1.9/7.46 ppm; editing pulse duration = 15 ms, TR = 2000 ms; 320 averages; 30 × 30 × 30 mm^3^; medial parietal lobe voxel).

We allocated 40 of 48 selected in-vivo subsets (15360 transients) to the training subset (12800 transients), and the rest of the subsets (2560 transients) were used as an unseen test subset.

### 1.4 Deep Model

#### 1.4.1 The DAE proposed for Deep Learning-based Peak Referencing (dCrR, dCrRF)

The DAE is a type of deep artificial neural network that is created to learn the coding of data in an unsupervised manner. The fundamental underlying concept of autoencoders is to use the input data as the target, i.e., attempting to reconstruct the input data in the output layer ^14^. Typically, a DAE consists of two parts, namely, an encoder and a decoder.

Figure 1 illustrates the most common architecture of a DAE. The encoder function *h* = *f*(*x*) maps the *n*-dimensional input vector (*x* ∈ *R*^*n*^) to the *n*^′^-dimensional latent vector (*h* ∈ *R*^*n*′^), while the decoder function *x* = *g*(*h*) aims to reconstruct the *n*-dimensional output vector (*x* ∈ *R*^*n*^) from the latent space representation. The mathematical expression of a DAE can be written as follows:

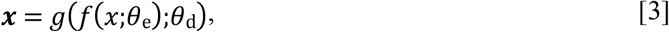

where *θ*_e_ and *θ*_d_ are the parameters set of encoder and decoder layers, respectively.

**Figure 1.**
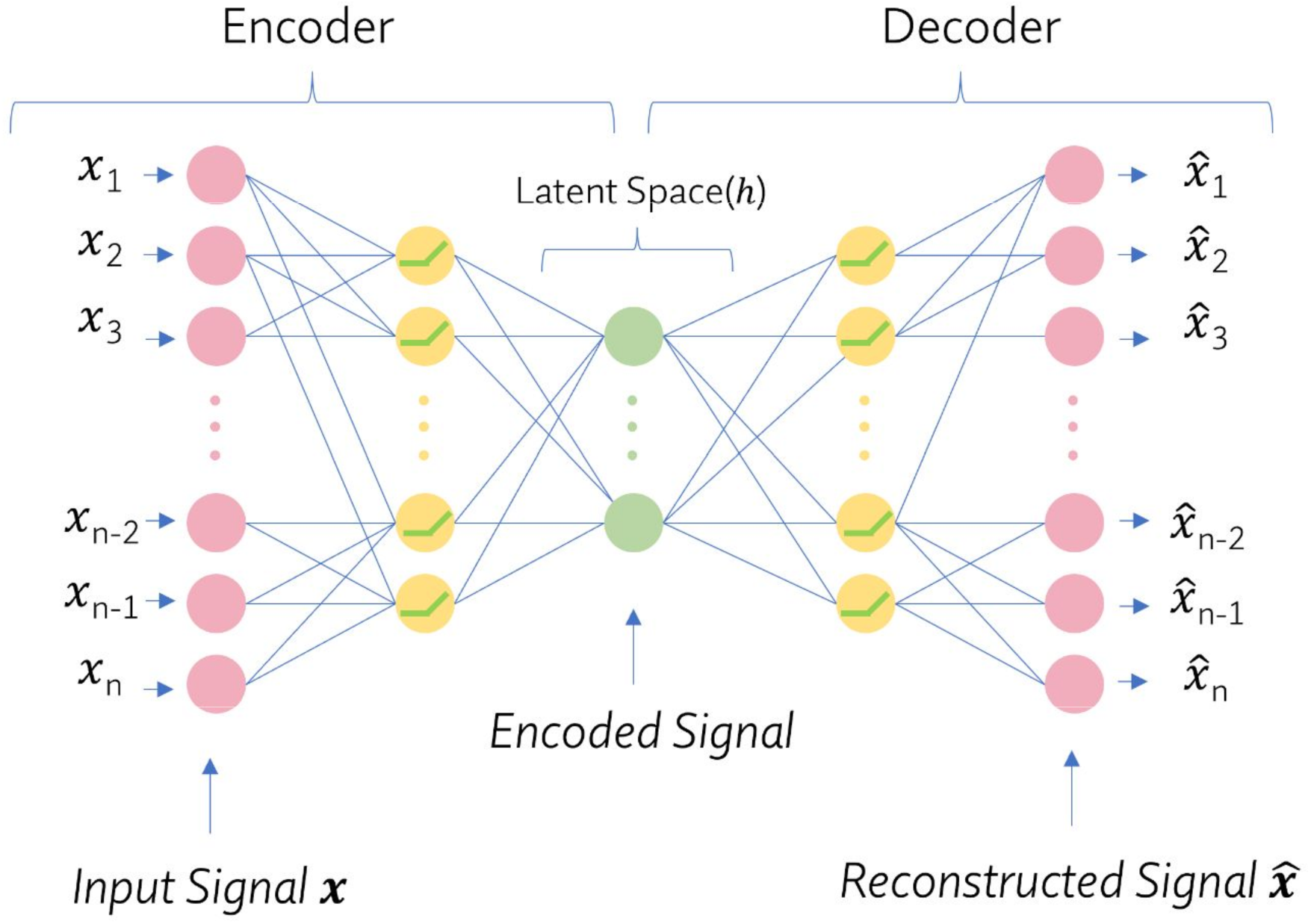
An illustration of a common DAE architecture with multiple non-linear hidden layers composed of ReLUs and FC layers to elicit non-linear features of the data. DAE, deep autoencoder, FC, fully connected; ReLU, rectified linear unit.

For FPC, the latent space representation must be interpretable parameters, such as frequency and phase shifts. Frequency and phase shifts can be estimated by fitting a certain metabolite peak to a model, e.g., a Lorentzian lineshape. Then, the frequency and phase shifts can be read from the model. To this end, we proposed a convolutional encoder / model-decoder ^16^ architecture. Our proposed DAE has a conventional encoder composed of a pipeline of a dropout layer ^33^, convolutional layers^34,35^, fully connected (FC) layers ^14,35^, and rectified linear unit (ReLu) layers^35^, which encodes a complex input signal into a latent space, and a decoder that reconstructs a Lorentzian lineshape of a certain peak in the input signal using the latent space parameters. The proposed DAE architecture is depicted in Figure 2. Since it has been shown that using complex-valued data improves the robustness and efficiency of fitting MRS data ^36^, the input and output of the proposed DAE were set to be complex signals in the time domain.

**Figure 2.**
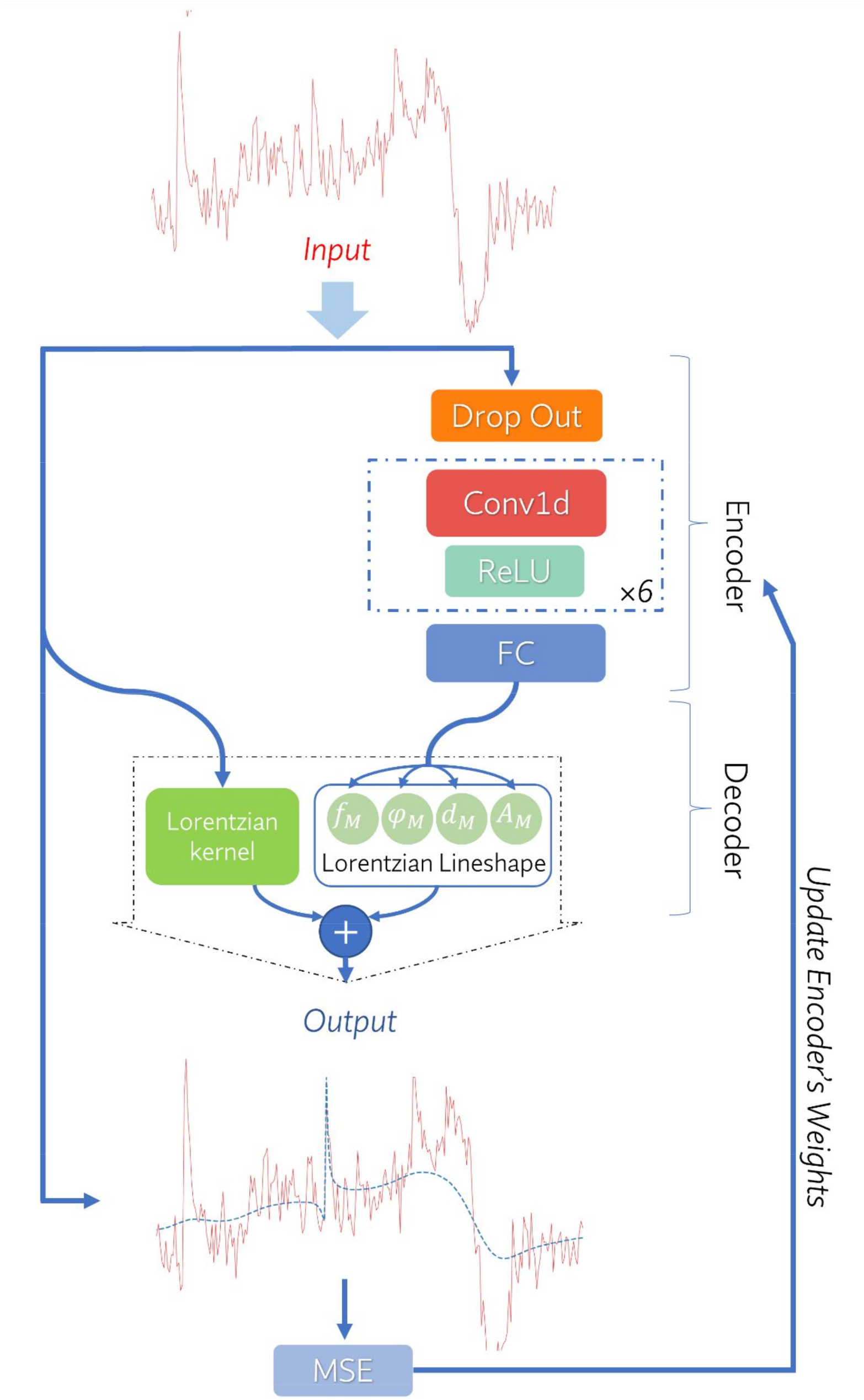
Illustration of the proposed convolutional encoder/model-decoder for dCrR method. The network’s input is a complex signal in the time domain, which is fed to the encoder. The encoder consists of a dropout layer, six convolutional blocks, and an FC layer (see details in Supplementary Information). A convolutional block (dashed square) is composed of a 1D convolution (Conv1d) layer followed by a ReLU layer. The model-decoder of the deep learning-based Creatine referencing (dCrR) (Eq. 5) reconstructs the output signal. The DAE was trained to encode the input vector in the time domain into parameters that can be used to reconstruct the output vector in the time domain. The proposed network is trained by minimizing the mean square error (MSE) between the input and the output. The input and output signals are depicted in the frequency domain for the sake of easier understanding. *A*_M_, *d*_M_, *f*_M_ and *φ*_M_ are the amplitude, the damping factor, the resonance frequency, and the zero-order phase of the Lorentzian lineshape, respectively. dCrR, deep learning-based Creatine referencing; FC, fully connected; ReLU, rectified linear unit.

Time-domain fitting of a single Lorentzian lineshape to a signal with several peaks using our proposed network is a challenging task in which the optimization algorithm aims to increase the linewidth to decrease the error. Previous studies ^3,12,29^ addressed these problems by fitting a signal in the frequency domain over a limited range and including a linear baseline in their model. We found that adding a rough estimate of a baseline, obtained by apodizing the input signal with Lorentzian kernel with a large linewidth, into the reconstruction function improves our fitting. Hence, the decoder part combines a mathematical model (Lorentzian lineshape) and the input signal, *x*, apodized with the Lorentzian kernel. The mathematical expression of the decoder can be written as follows:

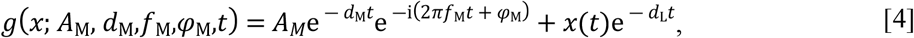

where *A*_M_, *d*_M_,*f*_M_ and *φ*_M_ are the amplitude, the damping factor, the resonance frequency, and the zero-order phase of the selected Lorentzian lineshape, respectively, and *d*_L_ is the linewidth of the Lorentzian kernel. Experimentally, *d*_L_ was set to 500 Hz in this study.

Training our proposed network is an unsupervised learning task that does not require ground truth frequency and phase shifts and can be done by minimizing the differences between the original input and the consequent reconstruction. In each iteration step of training, the parameters of the encoders are adjusted according to the gradient of the loss function with respect to the given parameters of the Lorentzian lineshape (*A*_M_, *d*_M_,*f*_M_, *φ*_M_).

In this study, Cr peak at 3.027 ppm was selected to be fitted by a Lorentzian line shape in the decoder. The FIDs were truncated to the initial 512 points for limiting the contribution of noise, which typically predominates in the later part of FIDs. Then, the truncated FIDs were utilized as inputs to the network. After training, the encoder of the proposed DAE (referred to as dCrR, deep learning-based Creatine referencing) was detached from the network and used to estimate the frequency and phase of the Cr peak in a test transient. Then the estimates were used for frequency and phase correction of the test transient. The pipeline of FPC of GABA-edited MEGA-PRESS transients is provided in Supplementary Information (Figure S6).

#### 1.4.2 The DAE proposed for Deep Learning-based Spectral Registration (dSR, dSRF)

With a simple modification, the proposed approach could estimate the relative frequency and phase shifts by fitting each signal to a reference signal and be applicable to various MRS experiments. In other words, the SR method could be employed in our proposed encoder / model-decoder network. This modification is referred to as deep learning-based spectral registration (dSR). For dSR, the mathematical expression of the decoder can be written as follows:

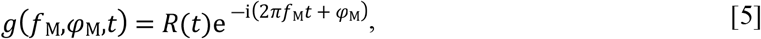

where *f*_M_ and *φ*_M_ are the frequency and the phase shifts of the input signal with respect to the reference scan *R*(*t*). Thus, in the Deep Learning-based Spectral Registration, the encoder estimates two parameters (relative phase and frequency shifts) instead of estimating the four parameters of a lineshape. The proposed DAE architecture for dSR is depicted in Figure S7. We proposed a new ML-based algorithm for finding the reference signal (see details in Supplementary Information).

#### 1.4.3 Training in the frequency domain over a limited frequency range

MRS signals may have unstable frequency components. Since all frequency components are present at all time points, unstable frequency components may bring errors into our proposed method. To avert this situation, the proposed architectures (dCrF and dSR) were trained over a limited frequency range (2.5 to 3.5 ppm) (referred to as dCrRF, deep learning-based Creatine referencing in the frequency domain, and dSRF, deep learning-based spectral registration in the frequency domain). Note that the input and the output of the network were still in the time domain.

### 1.5 Implementation Details and Training

All steps were run on a computer with a dual EPYC 7742 (2×64 cores) processor and one graphics processing unit (NVIDIA A100 40 GB). The DAE was implemented in Python with the help of the Pytorch lightning interface ^37,38^. The architecture of the network and training parameters were optimized using the Bayesian Optimization HyperBand algorithm ^39^ with the help of the Tune framework ^40^. The details of the optimization are given in Supplementary Information. All training was performed using the mean-squared error loss (MSE) and an Adam optimizer ^41^ with a batch size of 16, a learning rate of 4×10^−5^, and 150 epochs. The training progress for the simulated dataset is provided in the Supplementary Information (Figures S3 and S4).

An early-stopping strategy ^37^ was performed by monitoring the average error (Eq. 7) of the validation subset at the end of every epoch and stopping the training when no improvement was observed in 10 epochs. The SR and the SRF methods were tested using the FID-A toolbox^42^, the CrR and the CrRF methods with the Gannet toolbox^29^. All proposed methods (dCr, dCrF, dSR, and dSRF) were trained, validated, and tested using the simulated dataset, and the dCr method was trained and tested using the phantom and the big GABA datasets.

### 1.6 Statistics and Quality Evaluation

#### 1.6.1 Performance analysis

For the simulated dataset, in which the true shifts from the basis signal were known, the error was defined as the difference between the estimated and the true shifts. The accuracy and precision of an FPC method were established as the average and the standard deviation of the error, respectively. In addition, the performance of the dCrR method trained with the simulated dataset was investigated beyond the trained range of frequency and phase (−40 to 40 Hz and −180° to 180°, respectively).

For the phantom and the Big GABA dataset, where true shifts were not known, the quality of alignment was measured by comparing the similarity index (SI), which is the sum of all elements of a similarity matrix. Each element of the similarity matrix is the normalized scalar product (the equation as implemented is provided in the Supplementary Information) for each pair of spectra (FFT of a FID) over a limited frequency range (from 2.5 to 3.5 ppm). For the Big GABA dataset, similarity indices were compared using one-way analysis of variance (ANOVA) followed by the least significant difference post hoc tests to detect differences between dCrR-based correction, SR-based correction, and no correction. A *p*-value less than 0.05 was considered statistically significant.

#### 1.6.2 Performance against noise

The stability of the dCrR, dSR, and SR method against noise was evaluated. A set of transients was generated using the following procedure. First, a frequency shift of 5 Hz and a phase shift of 45° were added to *S*_basis_(*t*). Second, 20 realizations of a random complex Gaussian noise with a linearly increasing standard deviation were introduced to the shifted signal such that SNR was in the range of ∼8 to ∼110. The frequency and phase shifts of the generated set were estimated using the networks trained with the simulated dataset. FPC performance was evaluated as a function of SNR.

#### 1.6.3 Monte Carlo analysis

Monte Carlo (MC) studies were carried out to investigate the performance of the dCrR, dSR, and SR methods. A set of transients was generated using the following procedure. First, a frequency shift of 5 Hz and a phase shift of 45°were added to *S*_basis_(*t*). Second, 256 realizations of a random complex Gaussian noise with the same standard deviation were introduced to the shifted signal such that SNR was approximately 15. The frequency and phase shifts of the generated set were estimated using the networks trained with the simulated dataset. FPC performance of dCrR, dSR, and SR methods was compared.

## Results

The training and processing time of each proposed network for the simulated dataset is listed in Table 2. Approximately, frequency and phase estimation of one transient requires ∼3 ms. The evaluation of the performance of different methods (precision, SI, linewidth, the processing time of FPC, and training time for DAE) for the test signals of the simulated dataset and of the phantom are summarized in Tables 2, respectively.

**Table 2.** Comparison of our proposed method with existing commonly used FPC methods for the simulated dataset. Bold text indicates the best performance in each metric. SI of the test subset without FPC was 0.14. Precision: standard deviation of the difference between the estimated and the true shift, dSR, deep learning-based spectral registration; dSRF, dSR over a limited frequency range; SR, spectral registration; SRF, SR over a limited frequency range; dCrR, deep learning-based Creatine referencing; dCrRF, dCrR over a limited frequency range, CrR, Creatine referencing; Corr, frequency domain correlation, CorrF, Corr over a limited frequency range; SI, similarity index; Cr_linewidth_, the linewidth of Cr peak at 3 ppm; CPU, central processing unit; GPU, graphics processing unit.

|  |  | dSR | dSRF | SR | SRF | dCr | dCrF | Cr | Corr | CorrF |
| --- | --- | --- | --- | --- | --- | --- | --- | --- | --- | --- |
| Precision of frequency estimation (Hz) | All samples | 1.02 | **0.73** | 1.57 | 11.46 | 0.94 | 0.95 | 7.92 | 10.70 | 11.33 |
|  | Samples without nuisance peaks | 1.00 | **0.69** | 1.01 | 11.02 | 0.93 | 0.90 | 9.62 | 9.98 | 9.89 |
|  | Samples with nuisance peaks | 0.97 | **0.73** | 1.57 | 11.68 | 0.94 | 0.97 | 6.51 | 11.07 | 11.96 |
| Precision of phase estimation (°) | All samples | 5.87 | **4.89** | 7.44 | 15.92 | 6.62 | 6.79 | 32.34 | 16.87 | 17.46 |
|  | Samples without nuisance peaks | 4.16 | 3.34 | **2.61** | 16.75 | 6.89 | 7.00 | 38.80 | 16.62 | 16.46 |
|  | Samples with nuisance peaks | 5.75 | **5.52** | 8.50 | 15.34 | 6.35 | 6.52 | 26.68 | 17.02 | 18.07 |
| SI | All samples | 0.44 | **0.44** | 0.44 | 0.32 | 0.42 | 0.42 | 0.36 | 0.34 | 0.32 |
| $Cr_{linewidth}$ (Hz) | Averaged signal | 10.38 | 10.22 | 10.77 | 15.34 | **10.14** | 10.17 | 10.71 | 19.14 | 20.55 |
| Training Time (min) | All samples | **2.26** | 7.8 | - | - | 8.4 | 7.3 | - | - | - |
| Processing time (ms) CPU | Per signal | **3** | **3** | 57 | 71 | **3** | **3** | 53 | 33 | 29 |
| Processing time (ms) GPU | Per signal | **0.1** | **0.1** | - | - | 0.2 | 0.2 | - | - | - |

Figures 3a and 3b show scatter plots between resulting shifts estimated by dCrR, dCrRF and dCr against the true frequency and phase shifts. The highest precision (0.94 Hz and 6.62°) among the absolute FPC methods was achieved with the proposed dCrR. While dCrRF performed similarly to dCrR, CrR showed the lowest performance (7.92 Hz and 32.34°). All the proposed methods performed well in spectra with and without the simulated nuisance peaks. The agreement, estimated by R^2^ value, was high for dCrR (R^2^_frequency_ = 0.99 and R^2^_phase_=0.99) and dCrRF (R^2^_frequency_ = 0.99 and R^2^_phase_=0.99) and moderate for CrR (R^2^_frequency_ = 0.61 and R^2^_phase_=0.75). Figures 3c and 3d illustrate the results of dCrR on unseen test signals in which offsets are beyond trained bound (−40 to 40 Hz and −180° to 180°). The network showed poor performance beyond its trained bound (precisions for frequency and phase estimation were 8.69 Hz and 60.04°, respectively). Figures 3e and 3f show the spectra and the similarity matrix heatmaps obtained, respectively, before and after the FPC tested. The dCrR method increased the SI in the visualized test signals from 0.14 to 0.42.

**Figure 3.**
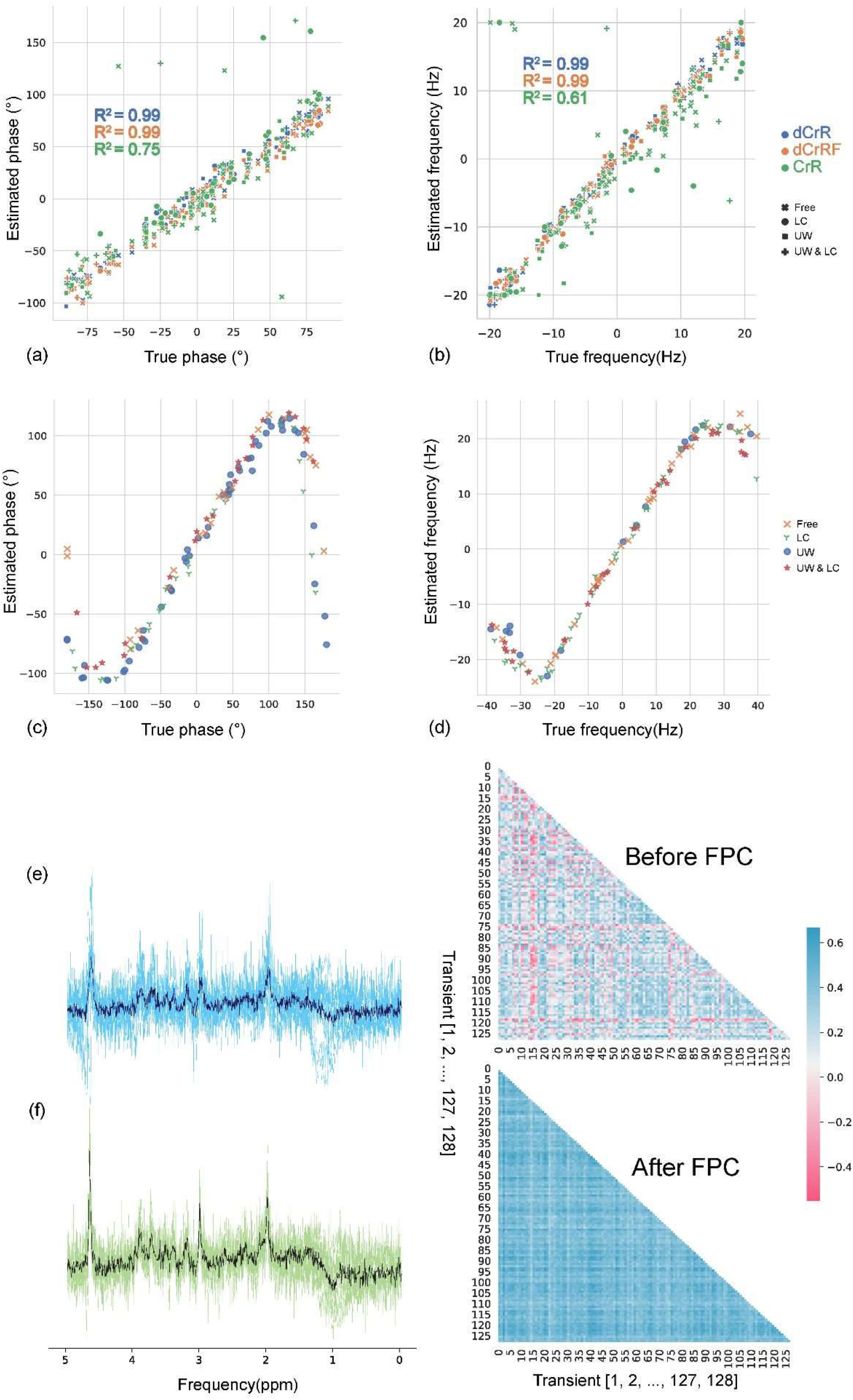
Testing results of the simulated dataset. (a,b) Plots of DL-estimated frequency (a) and phase (b) shifts against the actual shifts, respectively. R^2^ values of method types are color-coded. (c,d) Testing the dCrR method beyond the trained range of frequency and phase (−40 to 40 Hz and −180° to 180°, respectively). Plots of DL-estimated frequency (c) and phase (d) shifts against the actual shifts, respectively, using dCrR. (e) Uncorrected spectra and their similarity matrix. (f) The same spectra after FPC using dCrR and their similarity matrix. Dark blue and green spectra show the average uncorrected and corrected spectra, respectively. dCrR, deep learning-based Creatine referencing; dCrRF, dCrR over a limited frequency range, CrR, Creatine referencing; Free, without any nuisance peak; UW, with unstable water peak; LC, with Lipid peak.

Figure 4 shows a pairwise correlational comparison of the relative FPC methods (our proposed dSR and dSRF with SR and SRF). The R^2^ value of each method pair and the true value of shifts are reported in the corresponding axes. The R^2^ indicates that there is a high degree of agreement between the frequency and phase estimations between methods except for SRF. The agreement between the estimations of methods and true values (R^2^) was high for dSR, dSRF, and SR and was low for SRF (R^2^_frequency_ = 0.03 and R^2^_phase_=0.94).

**Figure 4.**
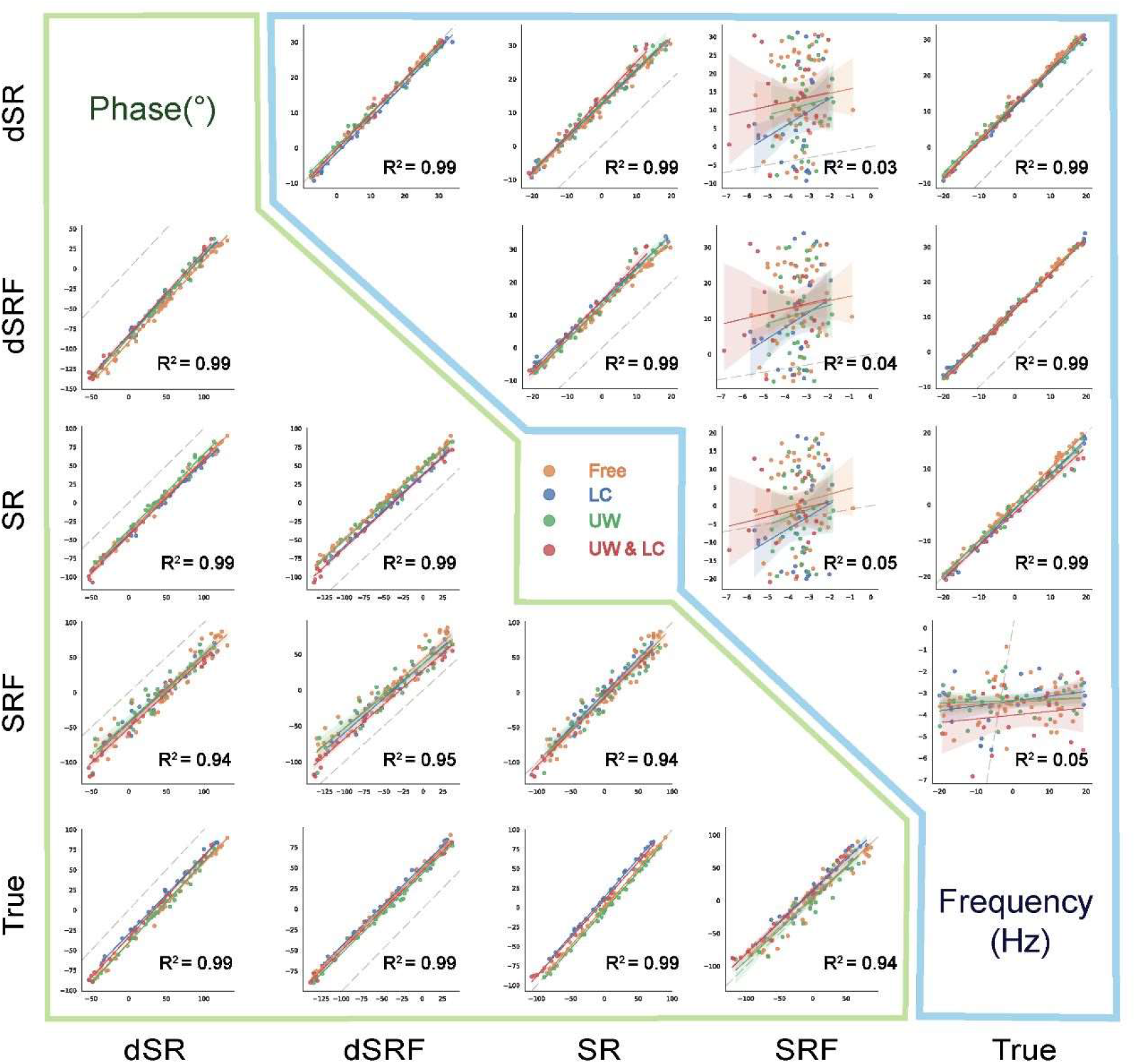
Pairwise comparison of the results of the SR-based FPC methods for the test subset of the simulated dataset. The upper triangular shows the correlations of frequency estimations, and the lower triangular shows the correlations of phase estimations. The fitted lines represent the linear regression model and the 95% confidence interval. The type of simulated nuisance peak is color-coded. The total R^2^ values are calculated along the corresponding axes. The dashed grey lines are identity lines. Note that the reference signals of relative FPC methods were different, which resulted in shifting their scatter plot from the identity line. dSR, deep learning-based spectral registration; dSRF, dSR over a limited frequency range; SR, spectral registration; SRF, SR over a limited frequency range; Free, without any nuisance peak; UW, with unstable water peak; LC, with Lipid peak.

Figure 5 shows spectra from the test signals of the phantom dataset and the corresponding heatmaps of similarity matrices before and after correction using dCrR method. The dCrR method achieved the highest performance and increased the similarity among transients from 0.50 to 0.66, and decreased the linewidth of Cr peak at 3 ppm in the averaged spectrum by 1.5 Hz (Table 3).

**Figure 5.**
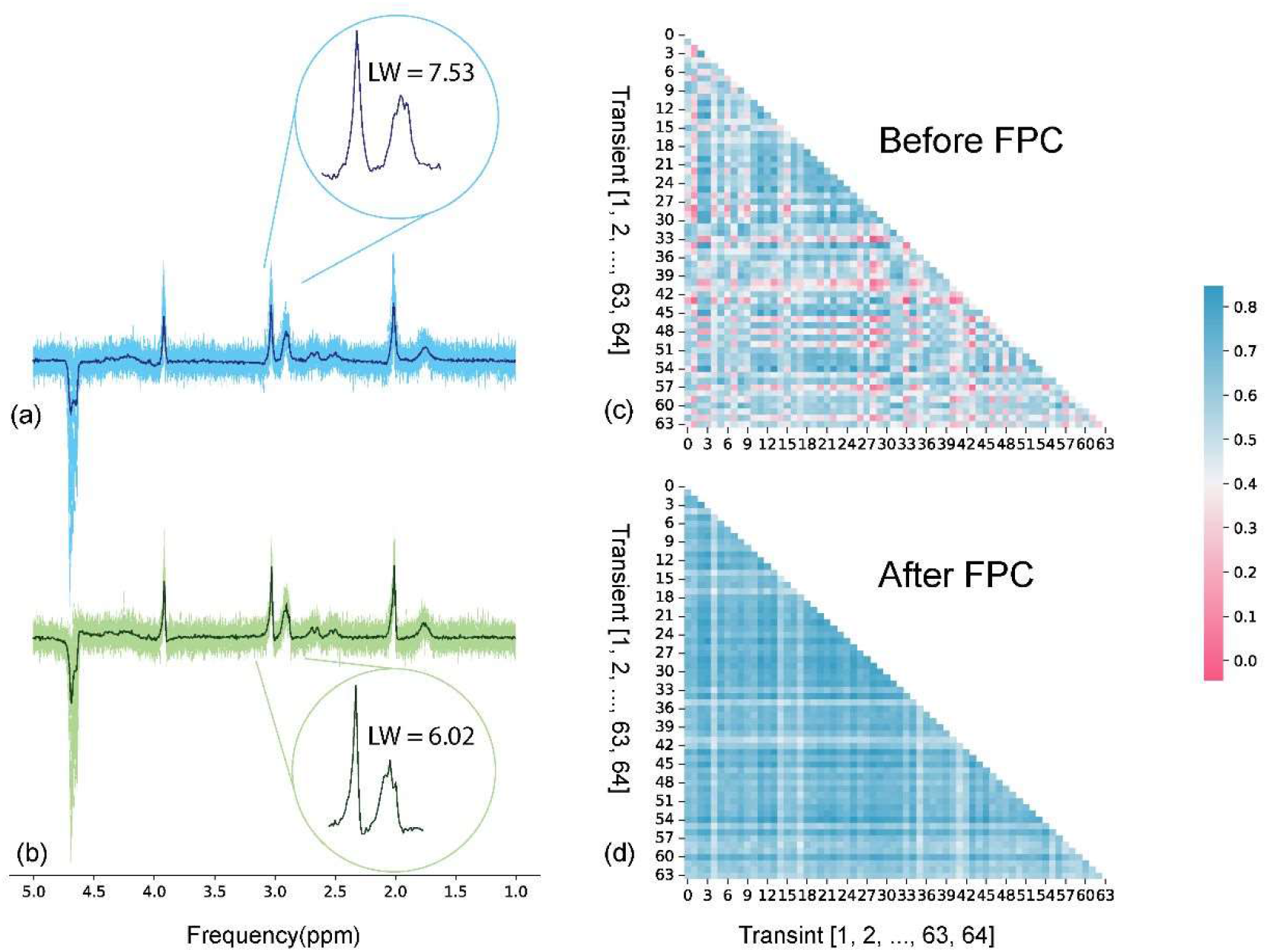
Frequency and phase correction of the phantom test subset using the dCrR method. Uncorrected (a) and corrected (b) spectra from the test subset. Dark blue and green spectra show the average uncorrected and corrected spectra, respectively. The circled inset shows the zoomed Cr peak at 3 ppm. The similarity matrix of 64 samples of the test subset before (c) and after (d) FPC. dCrR, deep learning-based Creatine referencing; LW, linewidth.

**Table 3.**
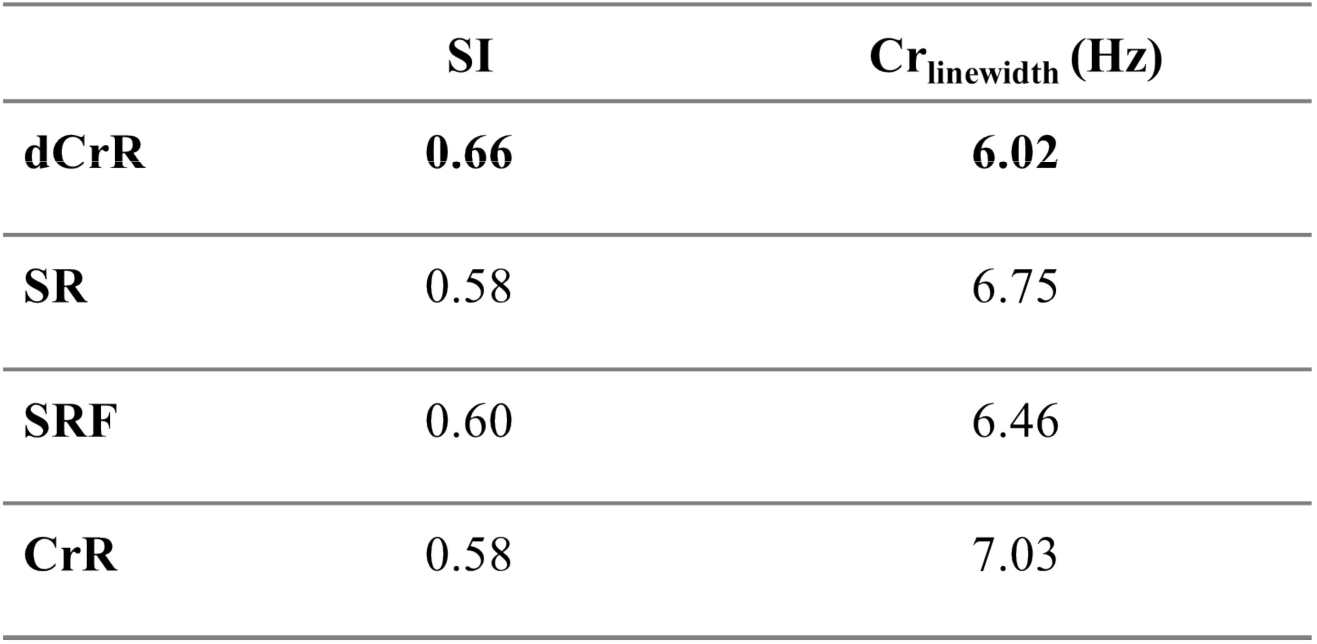
Comparison of our proposed method with existing commonly used FPC methods for the phantom dataset. SI of the test subset without FPC was 0.50. Bold text indicates the best performance in each metric. SR, spectral registration; SRF, SR over a limited frequency range; dCrR, deep learning-based Creatine referencing; dCrRF, dCrR over a limited frequency range, CrR, Creatine referencing; SI, similarity index; Cr_linewidth,_ the linewidth of Cr at 3 ppm.

The link between the absolute error in estimates and the SNR is shown in Figure 6 using a scatter plot, where the vertical and horizontal axes represent the absolute error of phase and frequency offsets estimation, respectively, and the size of the circles represents the SNR value of an individual signal. The comparison of the proposed dSR and dCrR with SR methods is presented. In low SNR, dSR and SR methods outperformed dCrR in terms of the precision of phase shift estimation, but dCrR demonstrated more resilience in terms of the precision of frequency estimation.

**Figure 6.**
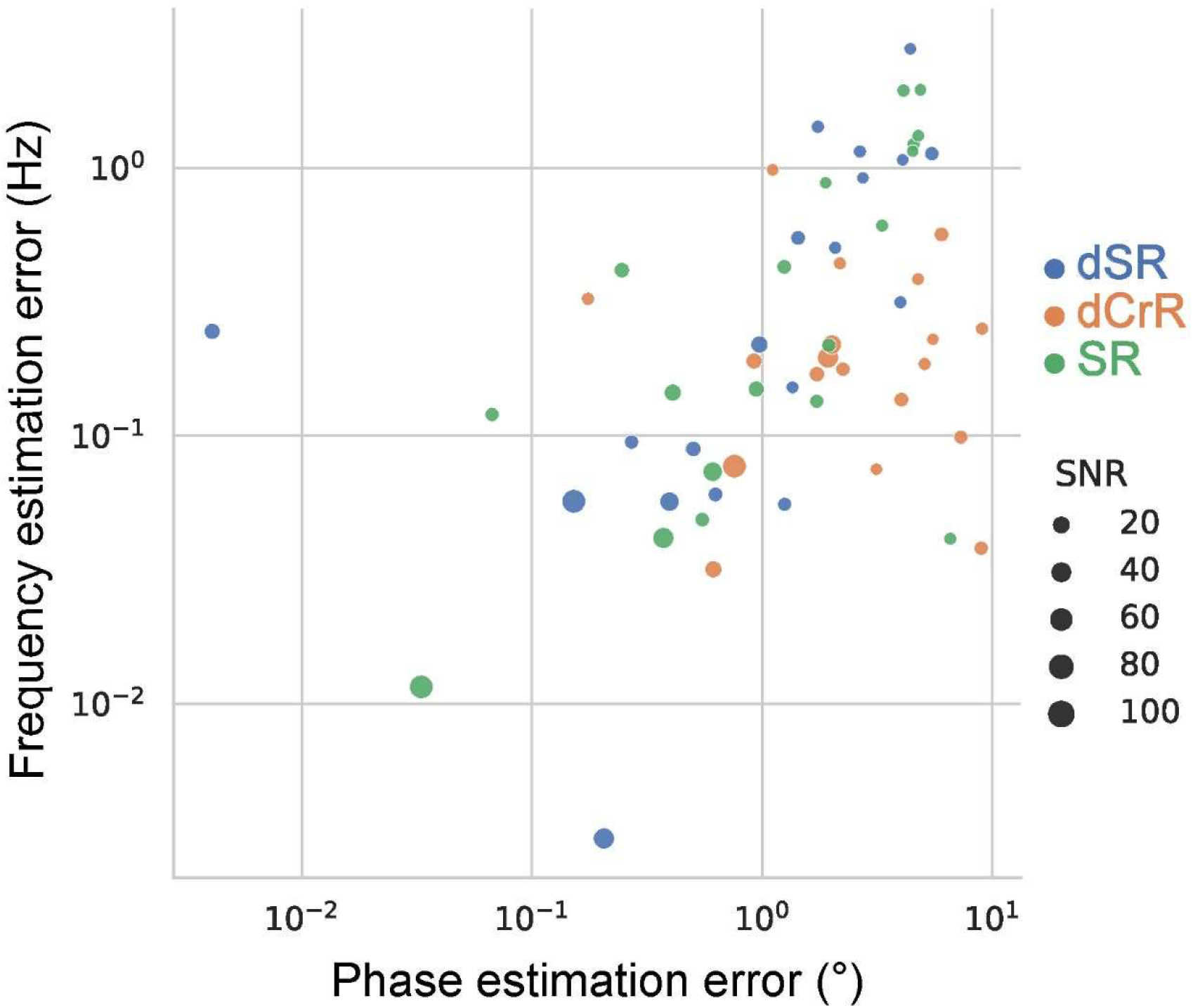
Comparison of frequency and phase correction precision of the dSR, dCrR, and SR methods over various SNR levels. The graph has logarithmic axes. dSR, deep learning-based spectral registration; dCrR, deep learning-based Creatine referencing; SR, spectral registration.

Figure 7 shows a comparison of the dSR, dCrR, and SR methods in the MC analysis using a scatter-plot visualization of the joint distribution of frequency and phase. For the simulated dataset, the mean error of dSR (1.36 ± 0.69 Hz and 0.51 ± 2.33°) and SR (−0.67 ± 0.81 and 3.15 ± 2.53°) showed similar performance, while dCrR performed a less precise in phase shift estimation (4.69 ± 0.77 Hz and −41.27° ± 5.827°). The true values of the frequency and the phase shifts were 5 Hz and 45°.

**Figure 7.**
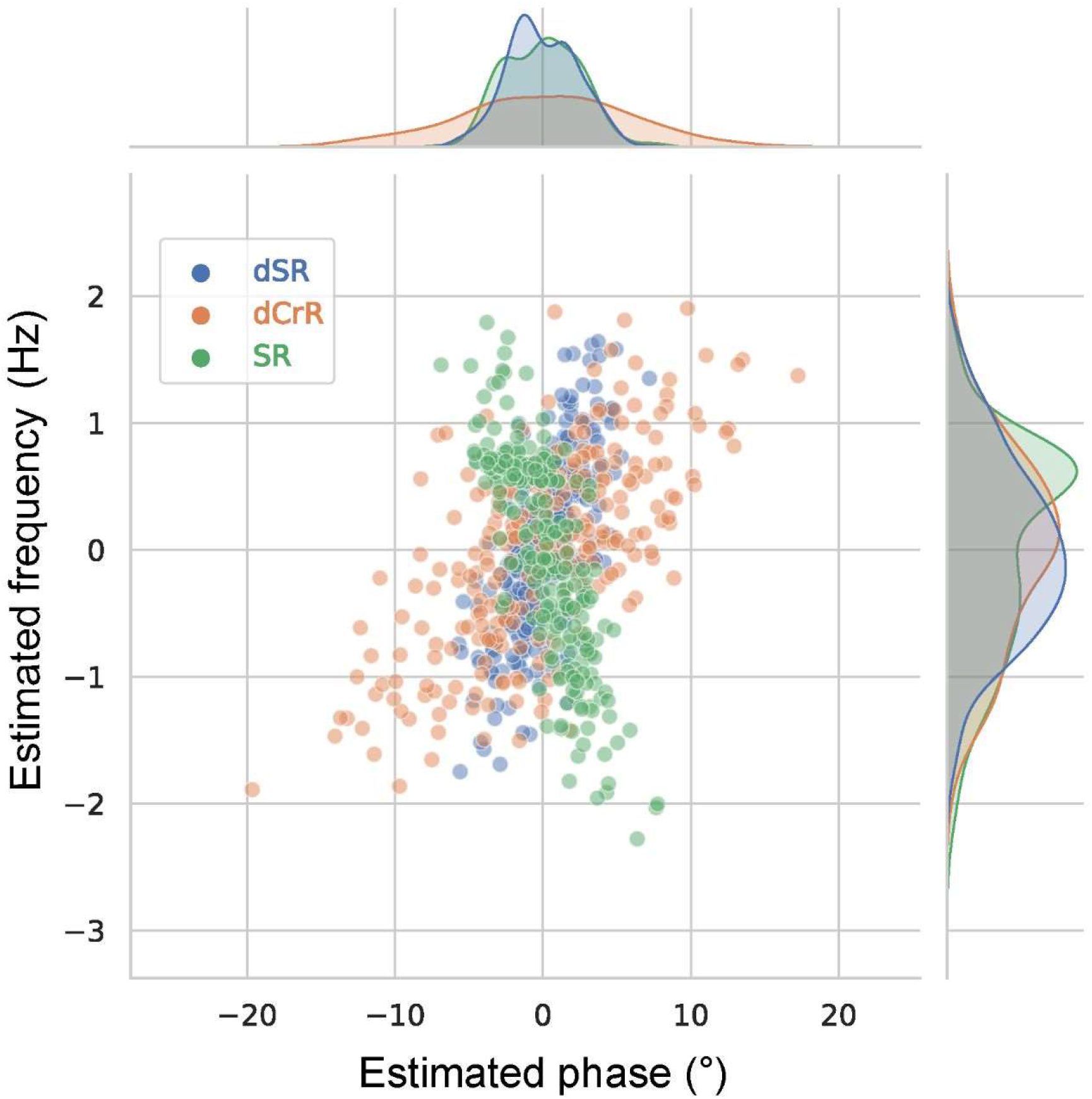
The results of MC analysis. Comparison of dSR, dCrR, and SR methods. For the sake of visualization of absolute and relative methods alike, the estimations from each method were subtracted from their average value. The result of dCrR method without subtracting from its average value can be found in Supplementary Information. dSR, deep learning-based spectral registration; dCrR, deep learning-based Creatine referencing; SR, spectral registration, MC, Monte Carlo.

Figure 8 illustrates an unseen test subset of GABA-edited in-vivo dataset (Site=1, Subject=3; 160 edited transients; 160 unedited transients) and a heatmap of their similarity matrix before and after FPC by the dCrR method. Our method increased the SI in the visualized test subset from 0.80 and 0.82 to 0.92 for edited (ON) and unedited (OFF) spectra, respectively. In all test subsets, the SI was increased from 0.88 ± 0.05 to 0.93 ± 0.02.

**Figure 8.**
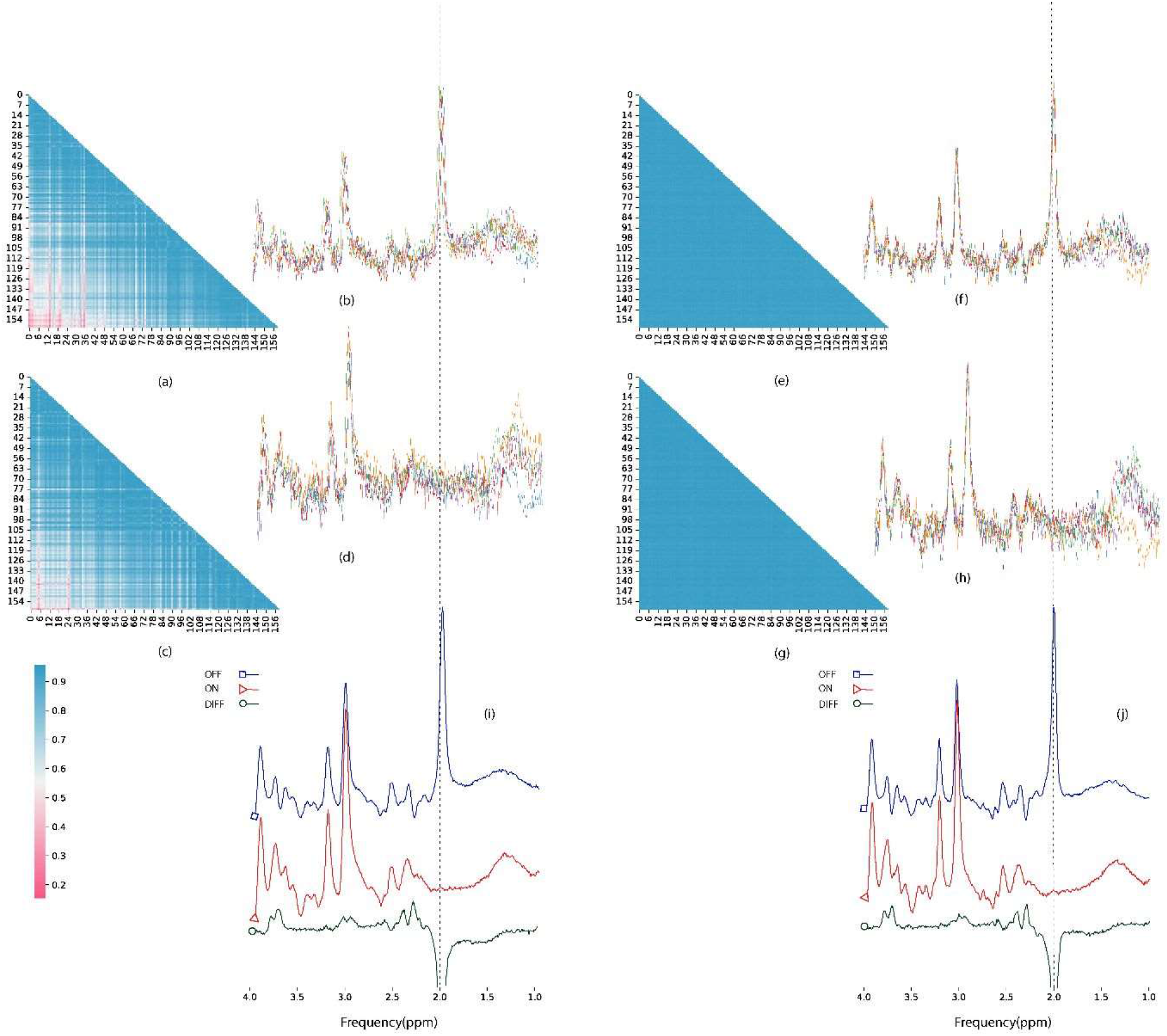
An example of FPC using dCrR for a test set in the GABA-edited in-vivo dataset. Unedited spectra (a) and their similarity matrix (b) before FPC. Edited spectra (c) and their similarity matrix (d) before FPC. Unedited spectra (e) and their similarity matrix (f) after FPC. Edited spectra (g) and their similarity matrix (h) after FPC. (i) Average uncorrected spectra (blue, unedited; red, edited) and their difference (dark green). (j) Average corrected spectra using dCrR (blue, unedited; red, edited) and their difference (dark green) dCrR, deep learning-based Creatine referencing.

Figure 9 shows the comparison of the results of dCrR-based and SR-based correction of ON and OFF transients of the test subsets. In all test signals, dCrR performed better than SR, as indicated by the mean SI (the mean SI was increased from 0.88 ± 0.05 [ON: 0.89 ± 0.04, OFF:0.87 ± 0.05] to 0.93 ± 0.02 [ON: 0.93 ± 0.02, OFF:0.93 ± 0.02] and 0.90 ± 0.04 [ON: 0.91 ± 0.03 OFF:0.89 ± 0.04] by dCrR-based correction and SR-based correction, respectively). The ANOVA test revealed differences between methods (*p* = 0.006). The post hoc tests showed differences between dCrR-based correction, dSR-based correction, and no-correction.

**Figure 9.**
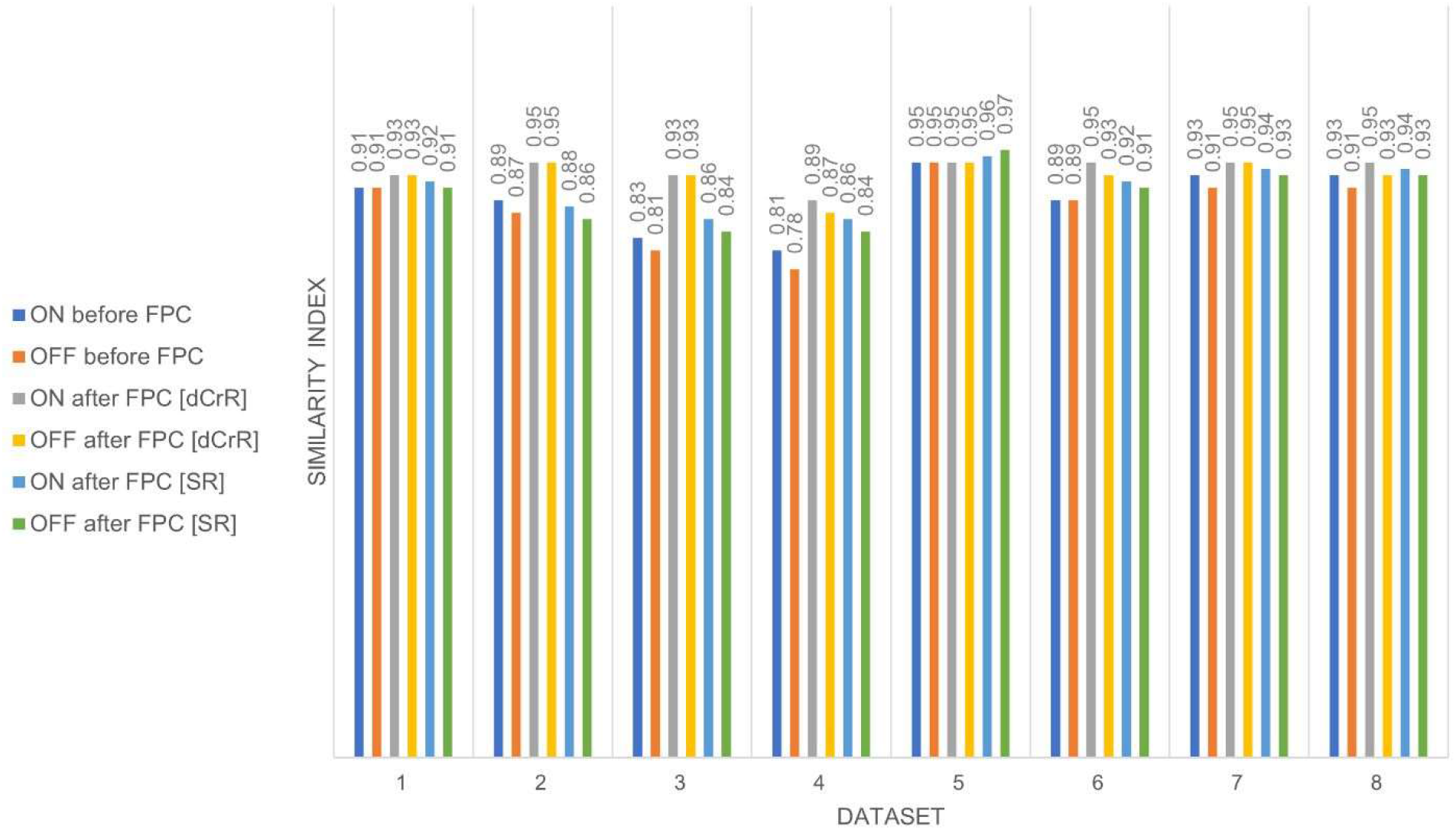
Similarity index comparing dCrR to SR for test signals of the Big GABA dataset. FPC, frequency and phase correction; dCrR, deep learning-based Creatine referencing; dSR, deep learning-based spectral registrtion.

## Discussion

In this study, the combination of DL and mathematical modeling was demonstrated to be able to provide FPC in simulated, phantom, and in vivo MRS data. We compared our method with five previously published methods: SR, SRF, Corr, CorrF, and CrR. We evaluated the ability of each of these methods to estimate the frequency and phase shifts in the simulated MRS dataset with known shifts and at varying SNR levels. Our results (Table 2) indicated an improvement in performance in terms of precision and the SI, the latter of which is a new measure proposed in this study for evaluating the FPC performance. Additionally, we compared our results to those obtained using other methodologies in terms of the linewidth of the Cr peak of the averaged signal obtained by summing corrected transients, and we discovered that the DL-based methodology performed comparably to others.

We found that the traditional FPC over a limited frequency range performed poorly in spectra with low SNR (Table 2), while our proposed FPC methods over a limited frequency range (dSRF and dCrRF) can perform equally (Table 2) to our proposed methods operating in the time domain (dCrR and dSR). Our methods also performed well in an MC study, where the phase- and frequency-shift estimate precisions were found to be very good and in general accordance with the SR method (Figure 7).

While the performance of conventional SR-based FPC methods can depend on the presence of nuisance peaks (Table 2 and ^3,7^), our result demonstrated that our methods functioned effectively in the simulated dataset regardless of the presence of nuisance peaks (Table 2).

Our proposed method can be trained in a few minutes (Table 2) because of using a one-dimensional signal as the input and a relatively tiny network. In the case of large MRS datasets or repeated measurements with the same conditions, the training time will be compensated by the FPC processing time, which is significantly shorter than in methods based on non-linear least squares (Table 2).

When the SNR of the test signals was lowered, the performance of our proposed methods (dCrR and dSR) and SR was reduced. When the SNR was decreased below the SNR of the training set, the performance deteriorated even further since the signal is dominated by noise. We observed that in low-SNR signals, dSR and SR methods performed better in phase shift estimation, while dCrR worked much better in frequency estimation (Fig 6).

Regardless of performance, the encoder / model-decoder provides a unique flexibility advantage. It leverages the underlying prior knowledge, which can be beneficial for estimating frequency and phase, independent of the kind of MRS data. Therefore, this approach may be used immediately to almost any sort of MRS data with little or no modification.

Contrary to the previous applications ^7,23^ of DL in FPC that used a supervised way utilizing simulated data, our proposed network was trained in an unsupervised way. This is advantageous since the majority of MRS data are unlabeled, and simulated datasets may not accurately reflect all in vivo circumstances, such as macromolecules and artifacts. Additionally, a single network was trained in this work to deliver both frequency and phase estimations by including prior knowledge into the decoder, while earlier work failed to train a single network.

It has been demonstrated that traditional FPC methods can benefit from additional information in a dataset; e.g., ^10^ addressed the problem of selecting a reference signal in SR by using a weighted average reference determined by mutual information in data. The encoder / model-decoder may assist in extracting patterns and information from data by introducing more complex models.

Overfitting is a common pitfall in DL ^14^. We implemented a dropout layer in the input to remove a part of the input randomly in every training step, which is a computationally inexpensive and very effective regularization strategy for decreasing overfitting and increasing the generalization of the network ^33^.

Training neural networks for regression problems necessitates a well-calibrated estimation. The results revealed a significant linear link between the true and estimated values (Figure 3 and Figure 4), indicating a well-calibrated estimation, although additional examination of the results is necessary.

Along with demonstrating the performance of the proposed approach on simulated and phantom data, the method was used to carry out FPC and enhance the similarity of signals in publicly accessible GABA-edited in-vivo MRS data. It should be emphasized that the proposed network was fed with both edited and unedited signals and trained simultaneously. The result shows the same performance for the edited and the unedited input.

One caveat in this study is that comparing processing time per transient between algorithms might not be widely valid since it might be affected by the parameters and conditions of algorithms. Utilizing the floating-point operations per second (FLOPS) to assess the computing cost ^43^ can help for a better comparison.

In the present study, we focused mainly on the validation of our proposed methods using in-silico ground truth knowledge and showed its application in GABA-edited in-vivo MRS data. We are aware that further evaluation and comparison with a more robust method ^10^ are needed.

## Conclusion

In general, our proposed time-domain FPC method, which is based on DL networks trained in an unsupervised way with complex data, may yield results comparable to previous FPC methods. The proposed approaches can perform absolute and relative FPC on extensively manipulated data in a shorter amount of time once the network is trained. Thus, our proposed approach could aid in the acceleration of analyzing large MRS datasets. Further study is needed to evaluate whether the same network could be trained and used for data originating in different studies.

## Acknowledgements

This work is part of the project that has received funding from the European Union s Horizon 2020 research and innovation program under the Marie Sklodowska-Curie grant agreement # 813120 (inspire-med). The authors thank Radim Kořínek, Ph.D. (Czech Academy of Sciences, Institute of Scientific Instruments, Czech Republic) for his valuable technical support.

## List of all Supplementary Information Captions

Supplemental Figure 1 (S1). Bin-based visualization of the result of the Monte Carlo analysis of the simulated dataset.

Supplemental Figure 2 (S2). Online monitoring of precision of a set of transients in the validation subset during training.

Supplemental Figure 3 (S3). Validation loss(MSE) versus training steps.

Supplemental Figure 4 (S4). Process flow of one-shot frequency and phase correction of J-difference edited MR spectra using a single deep neural network.

Supplemental Figure 5 (S5). Illustration of the proposed convolutional encoder-model decoder for dSR method.

Supplemental Table 1: Summary of the encoder s network.

Supplemental Text 1: Bayesian hyper-parameterization.

Supplemental Text 2: A novel ML-based algorithm for finding the reference scan for the deep Spectral Registration method.

Supplemental Equation 1: The equation for deriving each element of the similarity matrix.

## DATA AVAILABILITY STATEMENT

The source code is freely available at [https://github.com/amirshamaei/DeepFPC]. For questions, please contact the authors. For testing the Corr and CorrF methods, we developed a Matlab script, which is publicly accessible at [https://github.com/amirshamaei/Frequency-and-Phase-Correction-of-MRS-signals-Using-Cross-Correlation].

## Notes

https://github.com/amirshamaei/DeepFPC

### Competing Interest Statement

The authors have declared no competing interest.

