## SUPPLEMENTAL MATERIAL for "Model-Informed Unsupervised Deep Learning Approaches to Frequency and Phase Correction of MRS Signals"

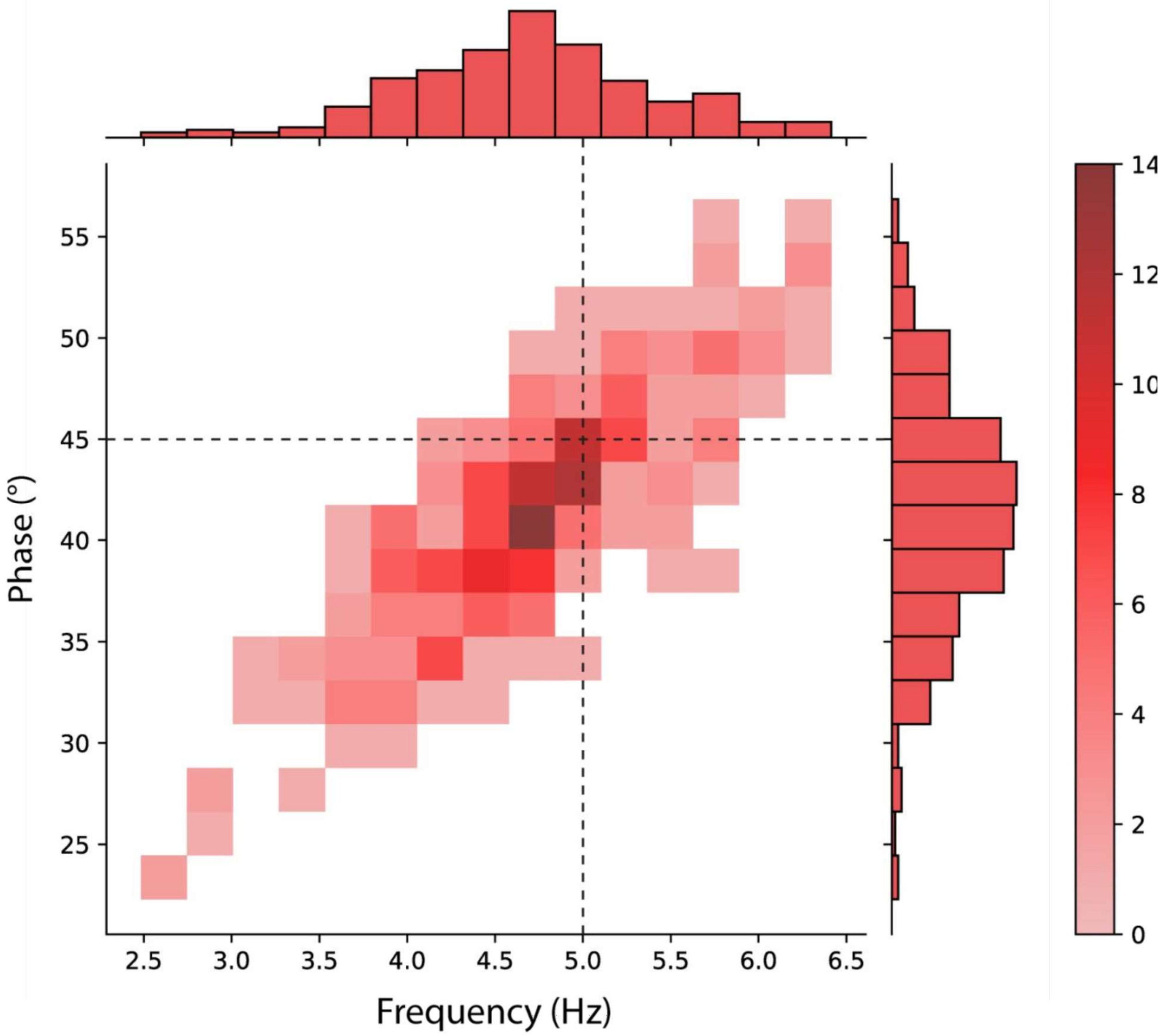


Supplemental Figure 1. Bin-based visualization of a joint distribution of frequency and phase estimation illustrates the result of the Monte Carlo analysis of the simulated dataset. Dashed horizontal and vertical lines show the true phase and frequency, respectively.

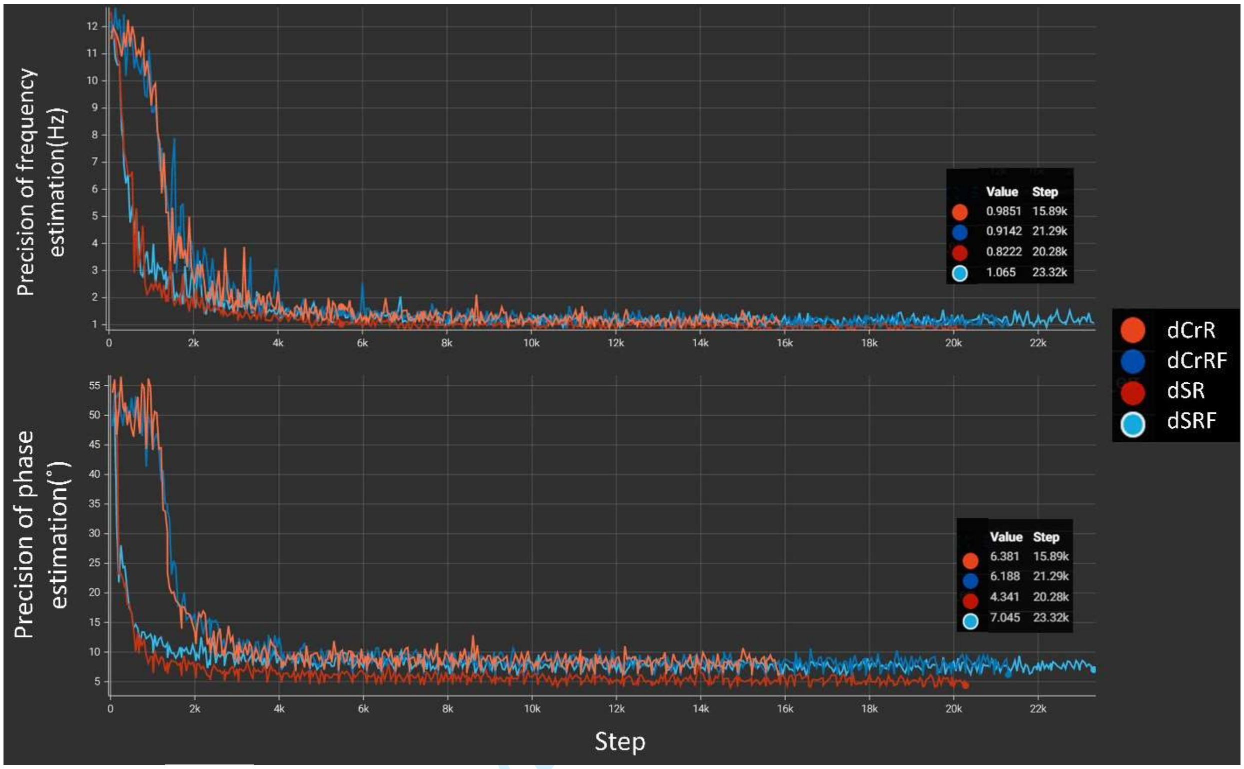


Supplemental Figure 2. Online monitoring of estimation precision during training for the simulated dataset(the validation subset). (top) the standard deviation of the error in predicted frequency shift versus training steps. (bottom) the standard deviation of the error in predicted phase shift versus training steps. Insets show the corresponding value at the last step of each method. dSR, deep learning-based spectral registration; dCrR, deep learning-based Creatine referencing; dCrRF, deep learning-based Creatine referencing in the frequency domain; dSRF, dSR over a limited frequency range.

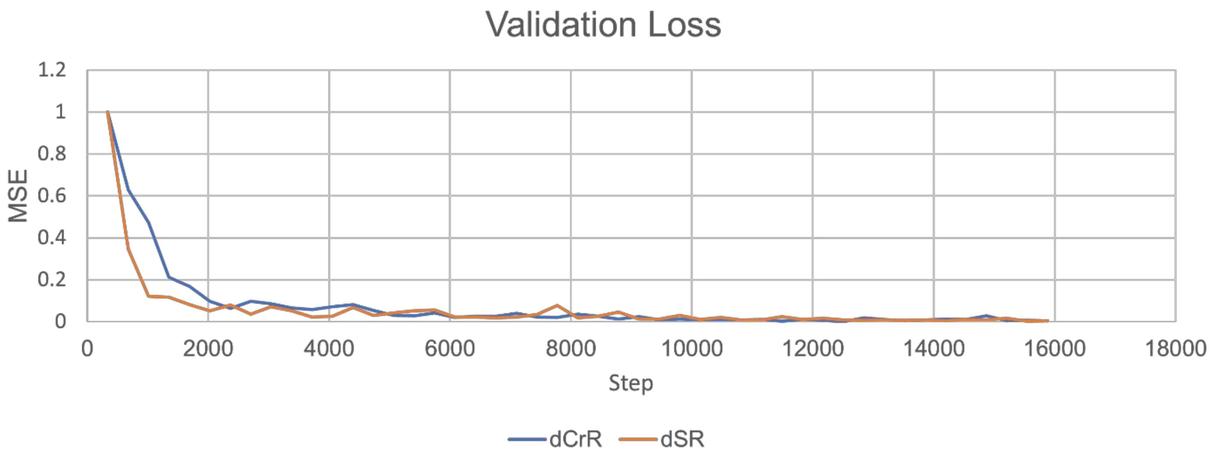


Supplemental Figure 3. Validation loss (MSE) versus training steps for the simulated dataset. dSR, deep learning-based spectral registration; dCrR, deep learning-based Creatine referencing; SR, spectral registration; MSE, mean square error.

For Peer Review

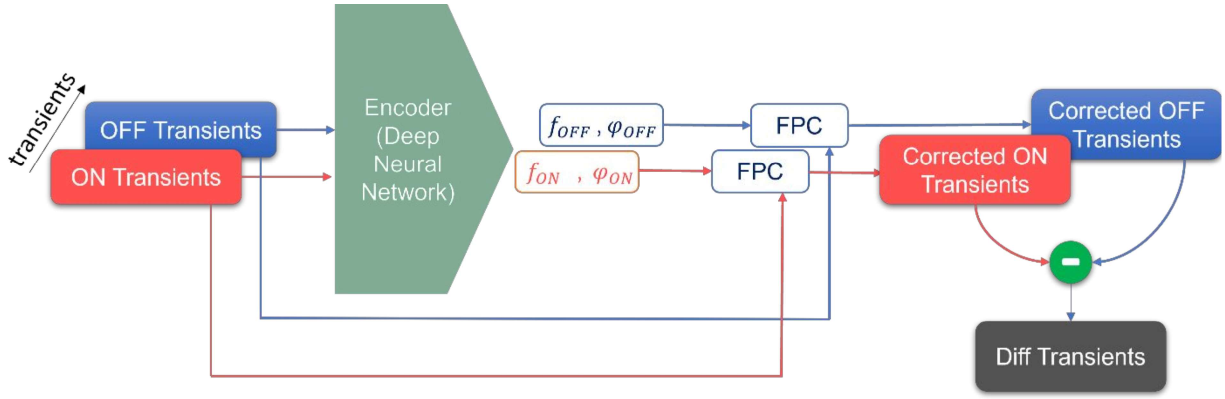


Supplemental Figure 4. Process flow of one-shot frequency and phase correction of J-difference edited MR spectra using a single deep neural network. Each input signal has two channels, one for the real part and another for the imaginary part in the time domain. ON and OFF transients are fed to the encoder concurrently, and the encoder estimates the frequency ($f_{ON}$and $f_{OFF}$) and phase ($\varphi_{ON}$and $\varphi_{OFF}$) shifts for each transient. The estimated shifts are added to the ON and OFF transients to achieve corrected ON and OFF transients. The difference (Diff transients) was obtained by subtracting the corrected OFF transients from the ON transients.

For Peer Review

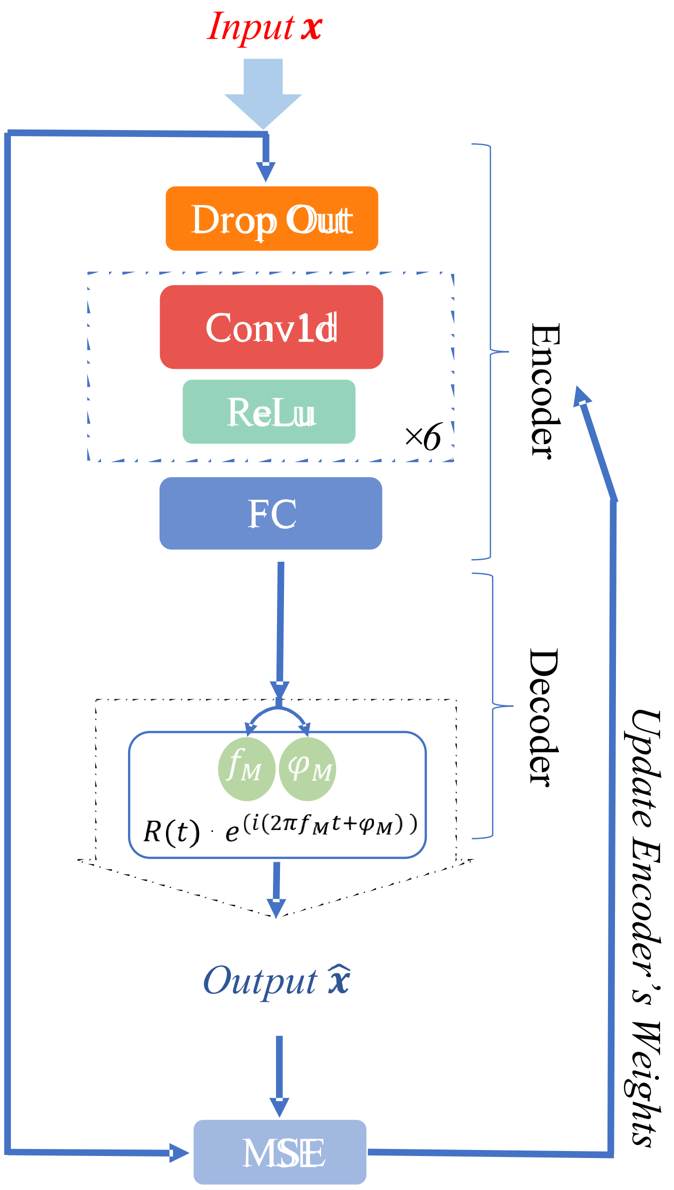


Supplemental Figure 5. Illustration of the proposed convolutional encoder–model decoder for the dSR method. The network's input is a complex signal ($\boldsymbol{x}$) in the time domain, which is fed to the encoder. The encoder consisted of a dropout layer, six convolutional blocks, and an FC layer (see details in Supplementary Information). A convolutional block (dashed square) is composed of a 1D convolution (Conv1d) layer followed by a ReLU layer. The model decoder of the deep learning-based spectral registration (dSR) (Eq. 5) reconstructs the output signal ($\boldsymbol{x}$). The DAE was trained to encode the input vector $\boldsymbol{x}$ in the time domain into parameters that can be used to reconstruct the output vector $\hat{\boldsymbol{x}}$ in the time domain. The proposed network is trained by minimizing the mean square error (MSE) between $\boldsymbol{x}$ and $\hat{\boldsymbol{x}}$. $f_M$ and $\varphi_M$ are the frequency and phase shifts of the input signal with respect to the reference scan. dSR, deep learning-based Spectral registration; FC, fully connected; ReLU, rectified linear unit; $R(t)$, reference signal.

Supplemental Table 1: Summary of the encoder's network. Conv1d, 1D convolution over an input signal; ReLU rectified linear unit; Flatten, reshaping it into a one-dimensional tensor; Linear, a linear transformation to the incoming data.

| Layer (type) | Output Shape [batch-size, channels, lentgh] | Number of Parameters |
|---|---|---|
| **Dropout** | [16, 2, 512] | |
| **Conv1d** | [16, 16, 255] | 176 |
| **ReLU** | [16, 16, 255] | |
| **Conv1d** | [16, 24, 127] | 1,176 |
| **ReLU** | [16, 24, 127] | 0 |
| **Conv1d** | [16, 32, 64] | 2,336 |
| **ReLU** | [16, 32, 64] | |
| **Conv1d** | [16, 40, 32] | 3,880 |
| **ReLU** | [16, 40, 32] | 0 |
| **Conv1d** | [16, 48, 16] | 5,808 |
| **ReLU** | [16, 48, 16] | 0 |
| **Conv1d** | [16, 56, 8] | 8,120 |
| **ReLU** | [16, 56, 8] | 0 |
| **Flatten** | [16, 1, 448] | 0 |
| **Linear** | [16, 1, 4] | 1,348 |

| | |
|---|---|
| **Total params** | 22,844 |
| **Trainable params** | 22,844 |
| **Non-trainable params** | 0 |

For Peer Review

Supplemental Text 1: Bayesian hyper-parameterization

The Bayesian Optimization HyperBand method(Falkner, Klein and Hutter, 2018) from the Tune framework(Liaw *et al.*, 2018) was utilized for the optimization of the encoder's hyperparameters. A total of 30 model trials were conducted, with the validation loss being monitored (MSE). The training was conducted for 50 epochs, and Tune's built-in early termination criteria were used(Liaw *et al.*, 2018).

Based on prior funding, we narrowed our parameters to only the four most significant parameters, which are as follows:

The learning rate: [1e-6, 1e-1]

The depth of the encoder, i.e., number of convolutional blocks: [2.. 7]

Using batch normalization layer: {Yes, No}

Batch size:[16.. 32]

Finally, the Bayesian Optimization HyperBand method explored the parameter space and identified the model with the best tuning and the least validation loss. The following table summarizes the tuned parameters:

The learning rate: 4.3235740152743133e-05

The depth of the encoder: 6

Using batch normalization layer: Yes

Batch size:16

Initially, the network architecture and training parameters were optimized in terms of MSE loss. However, since the network was trained in an unsupervised manner, the minimal MSE loss does not always correspond to the lowest frequency and phase estimation error. In the training of the simulated dataset, wherein the shifts were known, we monitored the $R^2$ (coefficient of determination) value of the estimated frequency- and phase-shifts during the training. We further tuned the parameters of the encoder to get the maximum $R^2$. Fine-tuned parameters are reported in the implementation details section.

For Peer Review

Supplemental Text 2: A novel Machine Learning-based algorithm for finding the reference scan for the deep Spectral Registration method

The K-means algorithm was used to cluster magnitude-valued signals in the training set. KMeans separated samples into two groups of equal variances by minimizing a criterion known as within-cluster sum-of-squares. The number of clusters was set to two, one for artifact-free transients and another for manipulated transients. The method was tested on the simulated dataset, and its accuracy (the proportion of correct predictions [both true positives and true negatives] among the total number of cases examined) was 76.1%.

The k-means algorithm divided transients into disjoint clusters, each described by the mean of the samples in the cluster. The means are commonly called the cluster "centroids" and reside in the same space.

K-means algorithm is an unsupervised method, so the clusters are not identified. Based on prior knowledge, contaminated transients have higher intensity in the initial points of their centroids. The first 10 points of each centroid were averaged, and the cluster with a lower average was identified as the cluster containing artifact-free transients. Then FIDs in the artifact-free cluster were Fourier transformed, and the transient with the highest SNR (highest peak of magnitude-valued signal between 2.5 to 4 ppm per the standard deviation of the real-valued noise) was selected as the reference scan. This method can be used for all SR-based methods.

For Peer Review

Supplemental Equation 1: The equation for deriving each element of the similarity matrix

$$SI = \frac{Re(S_i(f) \cdot S_j(f))}{\|S_i(f)\|\|S_j(f)\|}, \qquad [1]$$

where $S_i(f)$ and $S_j(f)$ are two complex vectors.

For Peer Review

Falkner, S., Klein, A. and Hutter, F. (2018) 'BOHB: Robust and Efficient Hyperparameter Optimization at Scale'.

Liaw, R. *et al.* (2018) 'Tune: A Research Platform for Distributed Model Selection and Training'. Available at: https://arxiv.org/abs/1807.05118v1 (Accessed: 25 January 2022).

For Peer Review
